# CTCF regulates hepatitis B virus cccDNA chromatin topology

**DOI:** 10.1101/2023.09.06.556185

**Authors:** Mihaela Olivia Dobrica, Christy Susan Varghese, James M Harris, Jack Ferguson, Andrea Magri, Roland Arnold, Csilla Várnai, Joanna L Parish, Jane A McKeating

## Abstract

Hepatitis B Virus (HBV) is a small DNA virus that replicates via an episomal covalently closed circular DNA (cccDNA) that serves as the transcriptional template for viral mRNAs. The host protein, CCCTC-binding factor (CTCF), is a key regulator of cellular transcription by maintaining epigenetic boundaries, nucleosome phasing, stabilisation of long-range chromatin loops and directing alternative exon splicing. We previously reported that CTCF binds two conserved motifs within Enhancer I of the HBV genome and represses viral transcripts, however, the underlying mechanisms were not identified. We show that CTCF depletion in cells harbouring cccDNA-like HBV molecules and in *de novo* infected cells resulted in an increase in spliced transcripts, which was most notable in the abundant SP1 spliced transcript. In contrast, depletion of CTCF in cell lines with integrated HBV DNA had no effect on the abundance of viral transcripts and in line with this observation there was limited evidence for CTCF binding to viral integrants, suggesting that CTCF-regulation of HBV transcription is specific to episomal cccDNA. Analysis of HBV chromatin topology by Assay for Transposase Accessibility/sequencing (ATAC-Seq) revealed an accessible region spanning Enhancers I and II and the basal core promoter (BCP). Mutating the CTCF binding sites within Enhancer I resulted in a dramatic rearrangement of chromatin accessibility where the open chromatin region was no longer detected, indicating loss of the phased nucleosome up- and down-stream of the HBV enhancer/BCP. These data demonstrate that CTCF functions to regulate HBV chromatin conformation and nucleosomal positioning in episomal maintained cccDNA, which has important consequences for HBV transcription regulation.

## INTRODUCTION

Hepatitis B virus (HBV), a hepatotropic DNA virus, can establish a chronic infection which is a global health burden affecting 296 million people worldwide resulting in 820,000 deaths each year due to associated cirrhosis and hepatocellular carcinoma (HCC) (World Health Organisation, 2019). HBV is a small, enveloped DNA virus [1] that enters hepatocytes upon binding to the sodium taurocholate co-transporting polypeptide (NTCP) receptor [2]. The viral nucleocapsid is uncoated and transported to the nucleus where the virus genome is released [3, 4]. The HBV genome, consisting of 3.2 kb partially double stranded relaxed circular DNA (rcDNA), is converted to a stable covalently closed circular form (cccDNA), which represents the template for six major viral transcripts with overlapping 3’ ends: pre-Core (pC) RNA (3.5 kb) encoding e antigen, pre-genomic (pg) RNA (3.5 kb) translated to polymerase and capsid antigen (core), preS1, preS2 and S RNAs (2.4, 2.1 kb) encoding the large (L), middle (M) and small (S) surface proteins (HBsAg) and X RNA (0.7 kb) translated to yield the X protein. The transcription of the viral RNAs is directed by four promoters that are regulated by two discrete but closely associated enhancer elements: Enhancer I activates the basal core promoter (BCP) which transcribes pg and pC RNAs, while Enhancer II activates Sp1, Sp2 and Xp promoters transcribing envelope and X RNAs [5, 6]. The pgRNA bound to the viral polymerase is encapsidated and replication occurs by reverse-transcription to form rcDNA. The resulting nucleocapsids are either recirculated to the nucleus or assembled into viral particles and further secreted [7]. Aberrant reverse-transcription events can generate a double stranded linear DNA (dslDNA) that can integrate into the host genome [7, 8]. HBV integrations and HBV-associated inter-chromosomal translocations have been identified in HCC-derived cell lines and liver biopsies from subjects with chronic hepatitis B (CHB) [9, 10]. Although integrated HBV DNA is replication-deficient, it is associated with HCC and represents a major source of HBsAg in CHB [8, 11].

Apart from the HBV canonical transcripts, spliced isoforms are produced and more than 16 variants of pC/pg RNA and 4 variants of preS2/S RNA have been identified [12]. Next generation sequencing data revealed the presence of spliced transcripts in experimental HBV cell culture models [13, 14], CHB liver biopsies [10] and serum from CHB patients [15] with SP1 being the most abundant. HBV splicing was reported to associate with HBeAg expression or HBsAg loss as a treatment outcome [10, 15] along with a poor response to interferon-α therapy [16], liver disease progression [17, 18] and risk of developing HCC [19]. The spliced RNAs have been reported to encode novel truncated or fusion proteins that may impact viral replication as demonstrated for SP1 [17, 20, 21], SP7 [22], SP10 [23] and SP13 [24].

HBV cccDNA represents one of the main therapeutic targets to achieve viral cure as it persists as a chromatinised episome in the nuclei of infected hepatocytes [25-27]. Current antiviral treatments consisting of nucleoside/nucleotide analogues (NUCs) suppress viral replication but do not eliminate cccDNA and despite the progression of liver disease being reduced, the risk for HCC remains high. Poor T- and B-cell immune response also contributes to cccDNA persistence in the liver [28]. HBV can reactivate following cessation of NUC therapy, rendering treatment as life-long [7]. In this context, attention is focused on identifying host factors responsible for cccDNA biogenesis and persistence. cccDNA is assembled into nucleosomes and the epigenetic status of the chromatinised cccDNA is essential for the coordinated transcription of the viral RNAs [27, 29]. Nucleosome mapping experiments have suggested nucleosome phasing up- and down-stream of the enhancers to maintain BCP accessibility and activity. In addition, the canonical +1 nucleosome, situated downstream of the BCP, aligns with RNA Polymerase II (RNA Pol II) enrichment in a similar manner to that observed in mammalian chromatin [27]. Although the phasing of nucleosomes within HBV cccDNA appears fundamental to the maintenance of transcriptional activity, the mechanisms governing the establishment and maintenance of nucleosome phasing are not known. We previously identified a role for the chromatin insulator CCCTC-binding factor (CTCF) to repress HBV transcription [30]. We demonstrated that CTCF binds two conserved motifs within Enhancer I (EnhI) and Xp regulatory elements, resulting in a repression of transcription from BCP and reduced pC/pg RNA levels. Here we provide evidence that direct association of CTCF with HBV cccDNA regulates chromatin accessibility at the viral enhancers and BCP. These data provide evidence that CTCF is important for the positioning of nucleosomes within the cccDNA and disruption of chromatin structure results in deregulated HBV transcription.

## METHODS

### Cell lines

HepG2-NTCP (obtained from Stefan Urban, Heidelberg University, Germany), HepG2-HBV-Epi (obtained from Ulrike Protzer, TUM, Germany), PLC/PRF5 (obtained from Mario Pirisi), Huh1 and Hep3B cells (obtained from Stephanie Roessler) and HepG2 2.2.15 cells [31] were grown in Dulbecco’s Modified Eagle’s Medium (DMEM) with Glutamax (Gibco) supplemented with 10% FBS (Gibco), 1X NEAA (Gibco) and 100 U/mL penicillin and 100 ug/mL streptomycin (Gibco). 500 or 380 μg/mL geneticin (Thermo Fisher) was added for propagating HepG2-HBV-Epi or HepG2.2.15 cells, respectively. The cells were seeded on collagen- (Sigma) coated plates and grown at 5% CO_2_, 37°C.

### Plasmids and cloning

The pUC57-1.3mer-HBV.D3 plasmid containing 1.3 length of HBV genome from D3 genotype (Peter Revill, Melbourne, Australia) was used as a template to generate pUC57-1.3mer-HBV.D3.CTCF.BS1-2 which contains previously described mutations in the CTCF binding sites within EnhI [30]. Two DNA sequences containing both mutated CTCF binding sites were synthesised (IDT) and ligated into pUC57-1.3mer-HBV.D3 following restriction digestion with *Kpn*I and *Mfe*I, or *Bsr*GI and *Pci*I, respectively. The resulting CTCF BS1-2 mutant plasmid was verified by sequencing (Source BioScience).

### siRNA transfection

HepG2-NTCP, HepG2-HBV-Epi, PLC/PRF5, Huh1 and Hep3B cells were transfected with either 25 nM CTCF-targeting or non-targeting control (scramble) smart pool siRNA (Dharmacon) using DharmaFECT4 (Dharmacon) as transfection reagent following the manufacturer’s protocol. The transfection mixture was removed after 24 h and cells were harvested 72 h post siRNA delivery.

### CHB liver biopsies

As previously reported, RNAs extracted from n=25 liver biopsies from CHB patients were used for quantification of HBV RNAs by qPCR [32]. Briefly, with informed consent, liver biopsy samples were collected and small sections were retained for research purposes that exceeded the requirements for pathological examination. These studies were approved by the local ethical committee (University of Oxford; CE90/19).

### HBV production and quantification

HBV stocks generated in HepAD38 cells were purified on heparin from the supernatants as described before [33]. HBV wild type (WT) and CTCF BS1-2mutant (CTCF BS1-2m) viruses were generated by transfecting HepG2-hNTCP cells with either pUC57-1.3mer-HBV.D3 WT or CTCF BS1-2m plasmid (10 μg plasmid/10 cm diameter dish) and cells maintained in media supplemented with 2.5% DMSO and 2.5% FBS. The supernatants were collected every four days for three weeks. The viral particles were precipitated with 10% PEG8000 overnight and separated by centrifugation at 4600 rpm for 1 h, at 4°C. The supernatant was discarded, and the pellet resuspended in DMEM to reach a 200X concentration. Samples of the viral concentrated stocks were treated with Turbo DNase (Invitrogen) following the manufacturer’s instructions. DNA was extracted with the QIAamp DNA kit (Qiagen) and quantified for HBV DNA levels by qPCR using a standard curve generated by serial dilution of HBV1.3 plasmid and HBV specific primers (Table 1).

**Table 1.**
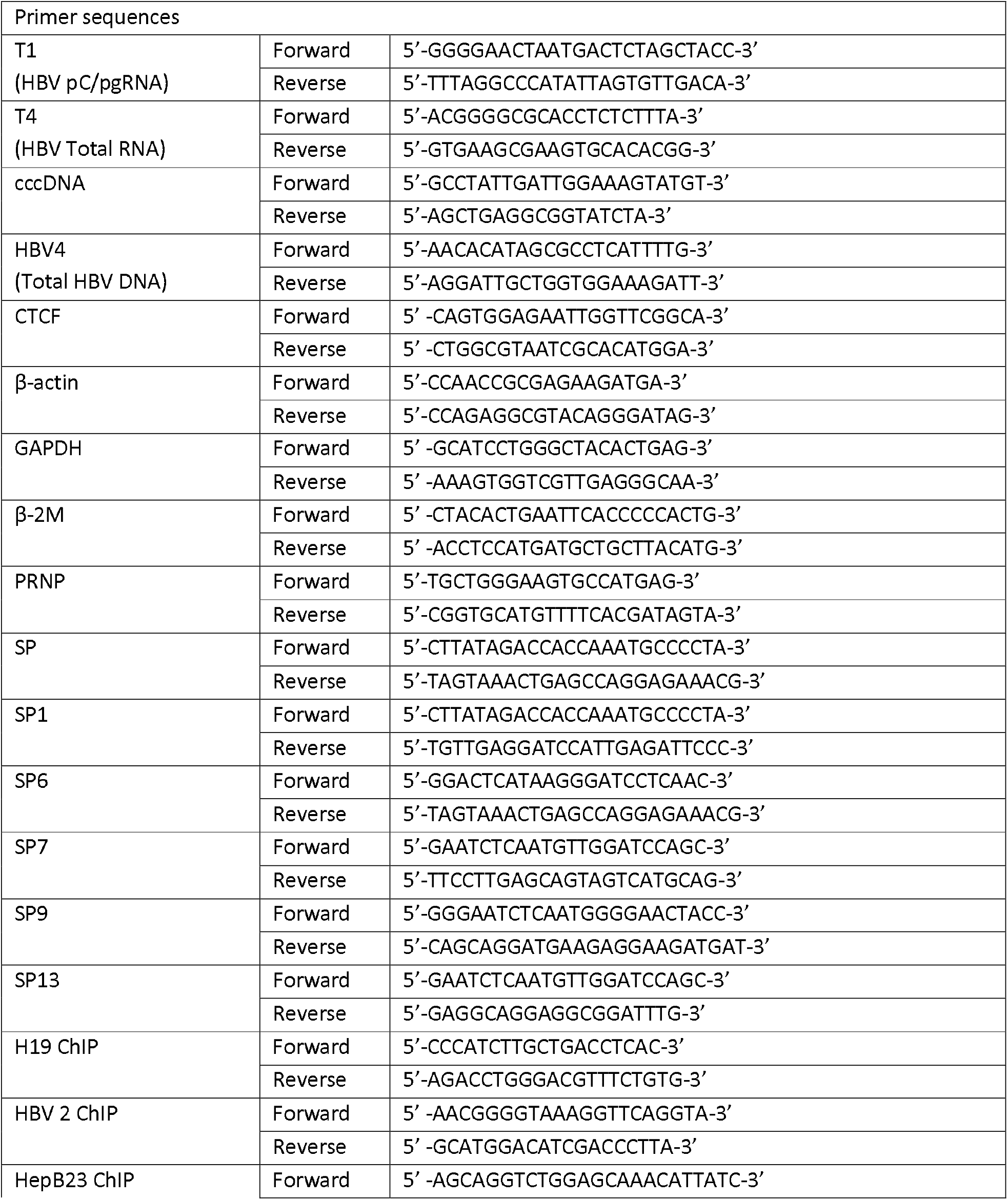
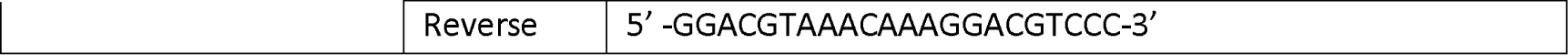
Oligonucleotide sequences used in this study where sequences are shown in a 5’-3’ orientation.

### HBV de novo infection

HepG2-NTCP cells were seeded on collagen-coated plates and infected with HBV purified by heparin affinity chromatography or PEG precipitated (WT or CTCF BS1-2m) at the indicated multiplicity of infection (MOI), in the presence of 4% PEG8000. The viral inoculum was removed after 16 h, the cells extensively washed with PBS and maintained in the presence or absence of 2.5% DMSO, as indicated. The supernatants were collected, and cells harvested at 3- and 6-days post infection (dpi). For the CTCF silencing experiments, cells were transfected with 25 nM scramble or CTCF siRNA at 3 dpi (as described above) and harvested after a further 72h (6 dpi). The supernatants were used to quantify HBeAg by ELISA and cells were lysed to extract RNA or DNA, as described below.

### RNA sequencing

HepG2-HBV-Epi cells were transfected with either 25 nM CTCF-targeting or non-targeting control (scramble) smart pool siRNA as described above. The cells were harvested 72 h later and RNA extracted (RNeasy mini kit, Qiagen). Libraries were prepared using Tru-Seq Stranded mRNA Library Prep kit for NeoPrep (Illumina, San Diego, USA) and 100 ng total RNA. Libraries were pooled and run as 75-cycle-paired end reads on a NextSeq 550 (Illumina) using a high-output flow cell. Sequencing reads were aligned to human (GRCh37) and HBV (HBV.NC_003977.2) genomes with STAR aligner (v2.5.2b) [34]. The analyses were performed on the CaStLeS infrastructure at the University of Birmingham.

### RNA extraction and RT-qPCR

RNA was extracted using the RNeasy kit (Qiagen) and DNase treated (Qiagen). RNA concentration was determined using Nanodrop 2000 spectrophotometer (Thermo Scientific). The RNA was subsequently used for reverse-transcription (RT) with the qPCRBIO cDNA synthesis kit (PCR Biosystems) and cDNA used for qPCR amplification of HBV transcripts using the qPCR SyBr green mix (PCR Biosystems) and specific primers (Table 1). GAPDH, β-Actin or β-2-Microglobulin were amplified as housekeeping controls as indicated.

### DNA extraction and qPCR

DNA was extracted using the genomic DNA kit (Qiagen) or the Allprep RNA/DNA kit (Qiagen). DNA concentration was determined using Nanodrop 2000 spectrophotometer (Thermo Scientific) and qPCR was further performed to detect HBV using the qPCR SyBr green mix (PCR Biosystems) and specific primers for total HBV DNA (Table 1). HBV-DNA levels were normalized to *PRNP* housekeeping gene. cccDNA was generated by treating the extracted DNA with T5 exonuclease (NEB) for 30 min at 37°C and measured by qPCR with specific cccDNA primers (Table 1).

### ATAC sequencing

HepG2-hNTCP cells infected with HBV wild-type or CTCF BS1-2m virus at MOI 400 were maintained in the presence of 2.5% DMSO. HBeAg levels were measured in the supernatants at 3 dpi and cells were harvested at 6 dpi. PLC/PRF5 cells were transfected with CTCF-targeting or non-targeting siRNA as described above and harvested at 3 days post-transfection. 100,000 cells were lysed, and total DNA tagmented and purified using the ATAC-seq kit (Active Motif) in accordance with the manufacturer’s protocol. The generated libraries were sequenced using Illumina NextSeq 550 paired-end sequencing. Reads were trimmed using Trimmomatic v0.39 with the parameters ILLUMINACLIP:TruSeq3-SE.fa:2:30:10 LEADING:3 TRAILING:3 SLIDINGWINDOW:4:20 MINLEN:50. Trimmed reads were aligned to HBV (HBV.NC_003977.2) using the BWA v0.7.17 aligner. Peaks were called within the aligned reads using MACS2 v2.2.7.1 with the parameters --keep-dup=auto. Statistical significance of differential DNA accessibility at EnhI/II/BCP was determined using a 1-sided t-test.

### HBeAg ELISA

Supernatants from HepG2-HBV-Epi cells or HepG2-hNTCP HBV-infected cells were collected at the indicated time points. HBeAg was detected using the HBeAg chemiluminescence immunoassay kit (Autobio). The luminescence signals were measured on a BMG Floustar Omega (BMG) and HBeAg was quantified using the standard curve provided by the manufacturer.

### SDS-PAGE and western blotting

Cells were lysed in urea lysis buffer (8 M urea, 150 mM NaCl, 20 mM Tris pH 7.5, 0.5 M β-mercaptoethanol) supplemented with protease inhibitor cocktail (Roche) and lysates were sonicated for 10 s at 20% amplitude using a Vibra-Cell sonicator (Sonics) fitted with a micro-probe. Total protein concentration was determined by BCA protein assay (Pierce). Proteins were denatured by heating at 95°C, separated by SDS-PAGE under reducing conditions and transferred to PVDF membranes (Amersham). The membranes were blocked in 5% milk and incubated with anti-CTCF antibody (Active Motif 61311; 1:1,000 dilution), anti-β-actin antibody (Sigma-Aldrich, 1:10,000 dilution) or GAPDH antibody (6C5, 1:1000 dilution) overnight at 4^⍰^C, washed and then incubated with HRP-conjugated secondary antibodies (1:10,000 dilution) for 1 h. The protein bands were visualised using the SuperSignal West Pico chemiluminescent substrate (Pierce) and images acquired on a G: Box mini (Syngene).

### Chromatin-immunoprecipitation (ChIP) and qPCR

PLC/PRF5 cells cultivated on 15 cm dishes were fixed with 1% formaldehyde for 10 min followed by 125 mM glycine treatment for 10 min. Cells were then washed with ice cold PBS, pelleted at 800 rpm, for 10 min and resuspended in lysis buffer (1% SDS, 10 mM EDTA, 50 mM Tris pH 8.1) supplemented with protease inhibitors (Roche) and incubated on ice for 30 min. Lysates were diluted 1:1 in ChIP Dilution Buffer (0.01% SDS, 1.1% Triton, 1.2 mM EDTA; 16.7 mM Tris pH8.1, 167 mM NaCl) and sonicated using a Biorupter (Diagenode) at high power for 15 min at 4°C (15s on/15s off cycles). Lysates were clarified by centrifugation (13,000 rpm, 10 min, 4°C), immunoprecipitated with either anti-CTCF (Active Motif 61311) or IgG control antibodies (agitation, overnight, 4°C) then incubated with Protein A-agarose beads (agitation, 1h 30min, 4°C). The pulled down samples were sequentially washed in low salt buffer (0.1% SDS, 1% Triton-X-100, 2 mM EDTA, 20 mM Tris pH8.1, 150 mM NaCl), high salt buffer (0.1% SDS, 1% Triton-X-100, 2 mM EDTA, 20 mM Tris pH 8.1, 500 mM NaCl), LiCl Buffer (1% Igepal, 1 mM EDTA, 10 mM Tris pH 8.1, 250 mM LiCl, 1% sodium deoxycholate) and twice in TE buffer (10 mM Tris pH8.0, 1 mM EDTA) before elution using elution buffer (0.1 M NaHCO3, 1% SDS). Eluates were incubated at 65°C overnight, 1400rpm, in the presence of 200 mM NaCl for reverse crosslinking of the complexes. Eluates were then treated with Proteinase K (45°C, 1400rpm, 4h) and RNaseA (37°C, 1400rpm, 30min) before cleanup using MiniElute PCR Purification kit (Qiagen). Purified DNA together with input samples before immunoprecipitation were amplified using qPCR SyBr green mix (PCR Biosystems) using two pairs of HBV primers: HBV2 and HepB23 (Table 1). Primers for human *H19* were used as a positive control for CTCF binding (Merck Millipore).

### PCR and DNA electrophoresis

DNA was extracted from HepG2, Huh1, PLC/PRF5, Hep3B cells and HBV control HepG2 2.2.15 cells and subjected to PCR amplification using the LongAmp Taq kit (NEB) and HBV primers spanning the CTCF binding motifs (HBV2 forward and Hep23B reverse). Products were separated and visualised by agarose gel electrophoresis.

### Statistics

Statistical analyses were performed using GraphPad Prism 9. The data are presented as mean +/- SD and Mann-Whitney U test was applied, unless otherwise indicated. Significance was considered for p<0.05 (*), p<0.01 (**), p<0.001 (***), p<0.0001 (****).

## RESULTS

### CTCF depletion increases spliced and non-spliced HBV transcripts

We previously reported that siRNA-mediated depletion of CTCF resulted in an increased abundance of HBV transcripts with no effect on cccDNA levels [30]. To further investigate the role of CTCF in regulating spliced and non-spliced HBV transcripts, CTCF was silenced in HepG2 cells containing episomal maintained HBV cccDNA-like molecules (HepG2-HBV-Epi; **Fig.1A**) and viral transcripts analysed by RNA sequencing. An average of 49.5 million reads were obtained per sample (range 37.5-66.5) with 93.5% (+/- 0.80 s.d.) uniquely mapped. Silencing CTCF resulted in a 1.7 fold increase in HBV sequences (reads per million, RPM) (p = <0.0001; **Fig.1B**). Although the number of HBV sequences were low (0.018% +/- 0.0045 s.d.), we were able to analyse the HBV transcriptome and identify splice events. Alignment of reads to the HBV genome sequence (NC_003977.2) showed an increase across the viral genome for the CTCF-depleted samples (**Fig.1C**). Alignment of HBV sequences to previously identified splice junctions [13] revealed a significant increase in splicing at the 2447^489 SP1 junction (p = <0.0001; **Fig.1D**) reflecting the predominance of this transcript as previously reported [13, 14]. Increased abundance of other transcripts spliced at 2447^2935 (SP7 or 13), 3018^489 (SP7, 14 or 18) or 3018^282 (pSP12) were noted, but these changes did not reach significance.

**Figure 1:**
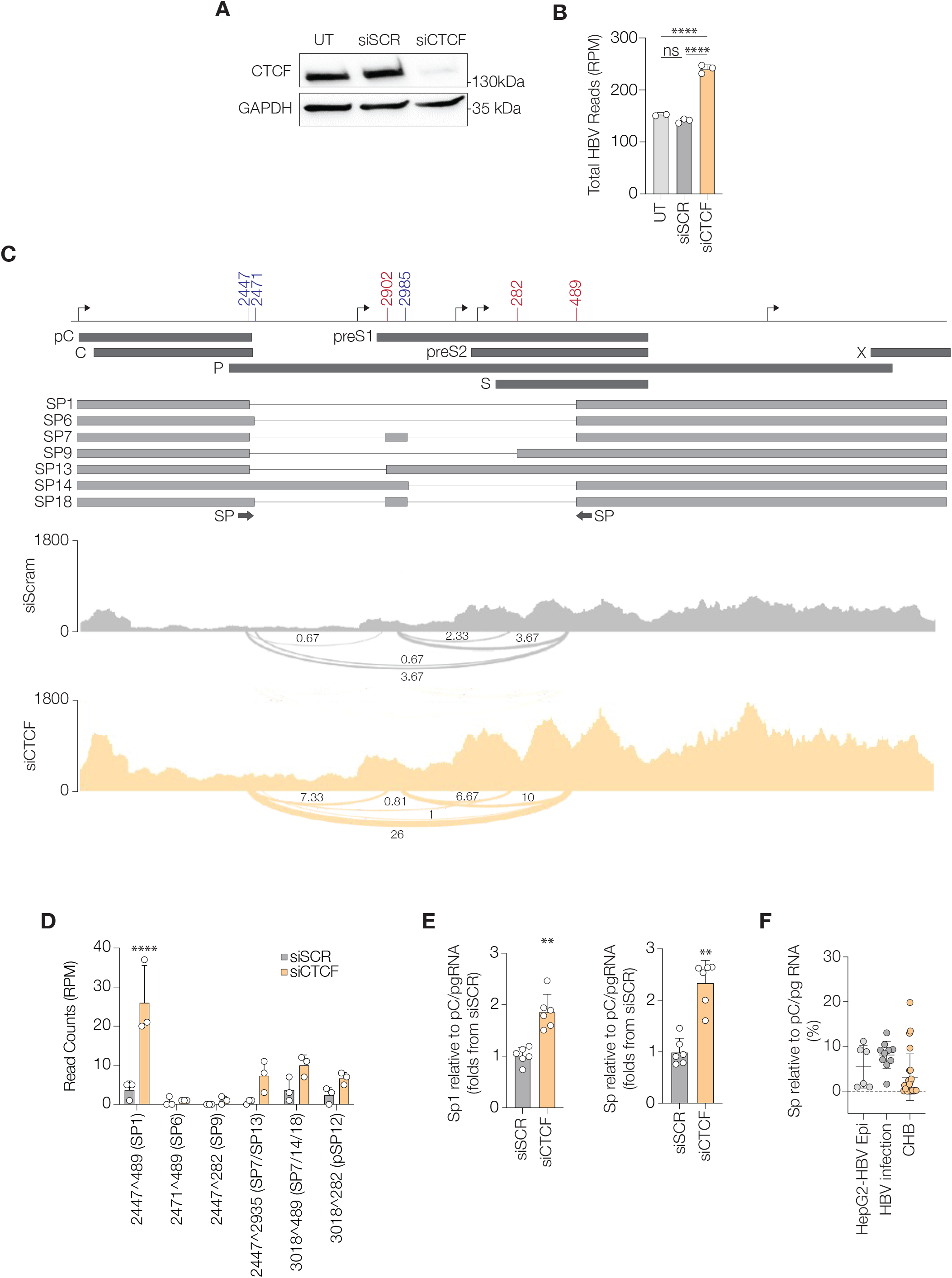
CTCF knockdown increases spliced and non-spliced HBV transcripts. HepG2-HBV-Epi cells were transfected with 25 nM scramble (siSCR) or CTCF siRNA (siCTCF) for 72 h or untransfected (UT). **(A)** CTCF protein was assessed by western blotting for CTCF and GAPDH. **(B)** PolyA+ selected mRNA was subject to short-read RNA-Seq and aligned to the HBV genome. The graph shows mean number of reads per million reads (RPM) that mapped to the HBV genome in triplicate samples of UT, siSCR and siCTCF +/- SD (**** p< 0.0001, one-way ANOVA with multiple comparisons). **(C)** Alignment of RNA-Seq data to the HBV genome, represented in a linear format. Open reading frames (dark grey) and spliced transcripts (light grey) are indicated along with annotation of TSS (black arrows) and splice donor (blue) and acceptor (red) coordinates. The mean RPM containing splice junctions is indicated on a representative alignment of siSCR and siCTCF samples. **(D)** Graph showing the mean +/- SD of RPM containing the indicated splice junction in three biological replicates (****p<0.000001, Student’s *t* test). **(E)** Expression of SP1 or total spliced HBV RNAs (SP) calculated by RT-qPCR relative to pC/pgRNA. Data show the mean fold change compared to siSCR +/- s.d. from three independent experiments performed in biological triplicates (**p<0.01, Mann-Whitney U test). **(F)** Expression of HBV SP calculated by RT-qPCR relative to pC/pgRNA in HepG2-HBV-Epi cells (n=6), HBV infected HepG2-hNTCP cells (n=14) or CHB liver biopsies (n=24).

To validate this increase in spliced transcripts following CTCF depletion in HepG2-HBV-Epi cells we analysed individual transcripts by RT-qPCR using primers designed to detect unique splice junctions on each of the transcripts (**Table 1**). Amplification of transcripts using primers specific for splice junctions 2447^489 (SP1), 2471^489 (SP6), 2449^2903-2987^490 (SP7), 2447^282 (SP9) and 2447^2935 (SP13) showed a significant increase in abundance relative to the cellular housekeeping transcript β-actin (**Supplementary Fig.1**). PCR detection of all spliced RNAs using the SP primers annotated in **Fig.1C** gave a similar profile to the SP1 amplicon, with increased spliced transcript levels in the CTCF silenced cells (**Supplementary Fig.1**). This increased abundance could reflect a general increase in pC/pg RNA or an altered regulation of specific splicing events. We therefore calculated the abundance of SP and SP1 relative to pC/pg RNA and noted a significant increase in the CTCF depleted cells (**Fig.1E**). The frequency of spliced transcripts in HepG2-HBV-Epi model was 5-15% of pC/pg RNA and we observed a similar ratio in *de novo* infected HepG2-NTCP cells or CHB liver biopsy samples (**Fig.1F**), consistent with our recent long-read sequence mapping of the HBV transcriptome [14]. Collectively, these data show a role for CTCF in repressing HBV RNA splicing.

### Differential roles for CTCF in regulating the abundance of spliced and non-spliced HBV transcripts

To extend our observations and confirm a role for CTCF in regulating the abundance of viral transcripts in *de novo* infection we inoculated HepG2-NTCP cells with HBV at an MOI of 250 or 500 genome equivalents (GE) per cell and silenced CTCF by siRNA transfection. Increased levels of pC/pg RNA, total RNA and SP1 along with HBeAg were detected in the silenced cells at both MOIs (**Fig.2A**). We previously identified two CTCF binding motifs in the HBV genome within EnhI (nt 1194-1209) and Xp (nt 1275-1291) [30]. We demonstrated that mutation of these sites alone or in combination reduced CTCF binding to cccDNA-like HBV minicircles (mcDNA) that was accompanied by an increase in HBV transcript abundance, indicating that CTCF may directly regulate HBV transcription. To address whether CTCF binding to authentic cccDNA regulates spliced RNAs we generated a modified virus bearing mutations of both CTCF binding sites (HBV CTCF BS1-2m) and infected HepG2-NTCP cells. The kinetics of cccDNA production in *de novo* infected HepG2-hNTCP cells were assessed by qPCR at 3 and 6 dpi. Infection with both wild type (WT) and CTCF BS1-2m viruses resulted in detectable cccDNA with no-significant variation between samples (**Fig.2B**). HBV transcript levels were normalised to cccDNA in each sample to infer HBV transcription. Infection with HBV-CTCF BS1-2m revealed a 3-fold increase in pC/pgRNA and 2-fold increase in total HBV RNA abundance compared to WT virus at 3 dpi (p = <0.05; **Fig.2B**). Although the magnitude of total RNA was sustained at 6 dpi in the mutant virus, the relative abundance of pC/pgRNA was less than 2-fold greater in HBV-CTCF BS1-2m (**Fig.2B**). The increased levels of pC/pgRNA in the HBV-CTCF BS1-2m infected cells was concomitant with an increase in secreted HBeAg (**Fig.2B**). Importantly, there was no change in SP1 levels in the WT or mutant infected cells, suggesting that CTCF regulation of spliced viral RNAs is not regulated by direct binding to the viral genome.

**Figure 2.**
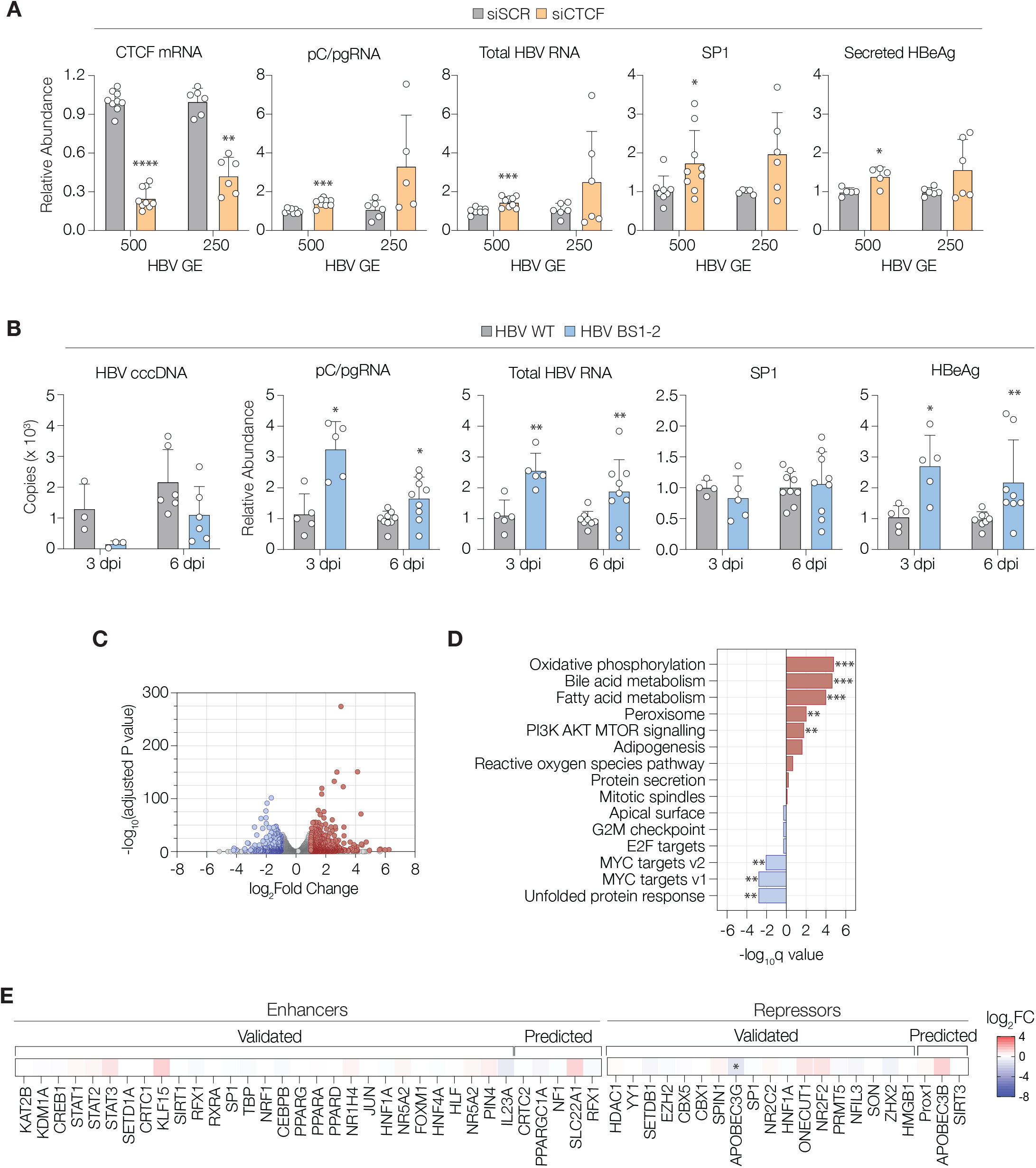
CTCF directly modulates abundance of HBV transcripts and indirectly modulates SP1. (A) HepG2-NTCP cells were infected with 250 or 500 HBV GE and at 72 h post-infection, the cells were transfected with 25 nM scramble (siSCR) or CTCF siRNA (siCTCF) and harvested after another 72h. RNA was extracted and CTCF and HBV transcripts quantifed by RT-qPCR. Secreted HBeAg was measured using the HBeAg chemiluminescence kit. Data for CTCF, pC/pg RNA transcripts, total HBV RNA and SP1 levels relative to pC/pg RNA and secreted HBeAg, respectively, are presented as fold change from siSCR controls (mean +/ -SD) from two independent experiments performed in biological triplicates. Statistical analysis was performed using Mann-Whitney U test for two group comparison (* p<0.05, ** p<0.01, *** p<0.001, **** p<0.0001). **(B)** HepG2-NTCP cells were infected with 400 GE HBV WT or virus harbouring CTCF binding sites mutations (HBV BS1-2m) in presence of 2.5% DMSO. Cells were harvested at 3 and 6 dpi and total DNA and RNA extracted. DNA was treated with T5 exonuclease and cccDNA detected by qPCR using specific primers. Data represents mean +/- SD of copy numbers of cccDNA adjusted to PRNP gene from two independent experiments performed in biological triplicates. pC/pg RNA, total RNA and SP1 levels relative to pC/pgRNA were detected by RT-qPCR. Secreted HBeAg levels were determined using the HBeAg chemiluminescence kit (*p<0.05,**p<0.01). **(C)** The impact of siCTCF on the host transcriptome. Differential gene expression was performed between siSCR- and siCTCF-transfected cells. Host gene expression was quantified and differential expression assessed by DESeq2. Genes with a log2FC of +/- 1 and an FDR <0.05 were deemed statistically significant. Upregulated transcripts are shown in red and downregulated in blue. **(D)** GSEA using the hallmarks gene sets to understand the pathways altered in siCTCF-transfected cells. Up- and down-regulated gene sets are plotted, (*q<0.05,** q<0.01, ***q<0.001). **(E)** DEGs were interrogated for the expression of factors previously reported to be enhancers or repressors of cccDNA activity (reviewed in [37]). Only APOBEC3G was significantly perturbed in the presence of siCTCF. Sequencing data presented in panels C-E were derived from n=3 replicates.

CTCF is a fundamental regulator of the cellular transcriptome and we hypothesised that silencing CTCF would dysregulate gene expression and perturb host gene expression that could influence HBV. We therefore interrogated our Illumina RNA-seq data derived from HepG2-HBV-pEpi cells and performed differential gene expression analysis between siSCR- and siCTCF-transfected cells. This revealed far reaching changes in gene expression with the upregulation of n=544 genes, and downregulation of n=331 genes (**Fig.2C**). Pathway analysis using the Hallmarks Gene Set from the Molecular signatures database [35] revealed the consequences of transcriptome perturbation, with genes implicated in oxidative phosphorylation, bile and fatty acid metabolism being amongst the most upregulated. Interestingly, the downregulated genes were associated with host cell processes such as the unfolded protein response, and MYC transcriptional activity (**Fig.2D**). As an obligate intracellular parasite, HBV is reliant on host factors to complete its replicative life cycle, and we analysed our differential gene expression analysis to assess whether CTCF regulated host factors which are known to regulate HBV [36, 37]. Whilst we saw a modest fluctuation in the expression of several host enhancers, we noted a significant reduction in the expression of apolipoprotein B mRNA editing enzyme, catalytic subunit 3G (APOBEC3G) [38]. Of note, APOBEC3G has been identified as a restriction factor of many viruses, including HIV and HCV, as well as HBV [39, 40].

### CTCF does not bind or regulate integrated HBV genomes

To assess whether CTCF can associate with integrated HBV DNA and regulate the abundance of viral transcripts in the context of the cellular genome, we depleted CTCF by siRNA transfection in cell models that carry integrated copies of HBV DNA: Huh1, PLC/PRF5 and Hep3B (**Fig.3A**). The levels of HBV-derived transcripts after CTCF silencing were measured by qRT-PCR. In all three cell lines tested, depletion of CTCF did not alter total HBV RNA levels (**Fig.3B**) even though the viral DNA around EnhI, which contains both CTCF binding sites, could be PCR amplified in all of the cell lines (**Fig.3C**). These data suggest that, in contrast to episomal maintained cccDNA, HBV integrants are not subject to CTCF-mediated transcriptional regulation. In support of this conclusion, there was no detectable enrichment of CTCF binding to viral integrants above the IgG control, as measured with two independent primer sets, despite robust immunoprecipitation of the cellular H19 positive control (**Fig.3D**).

**Figure 3.**
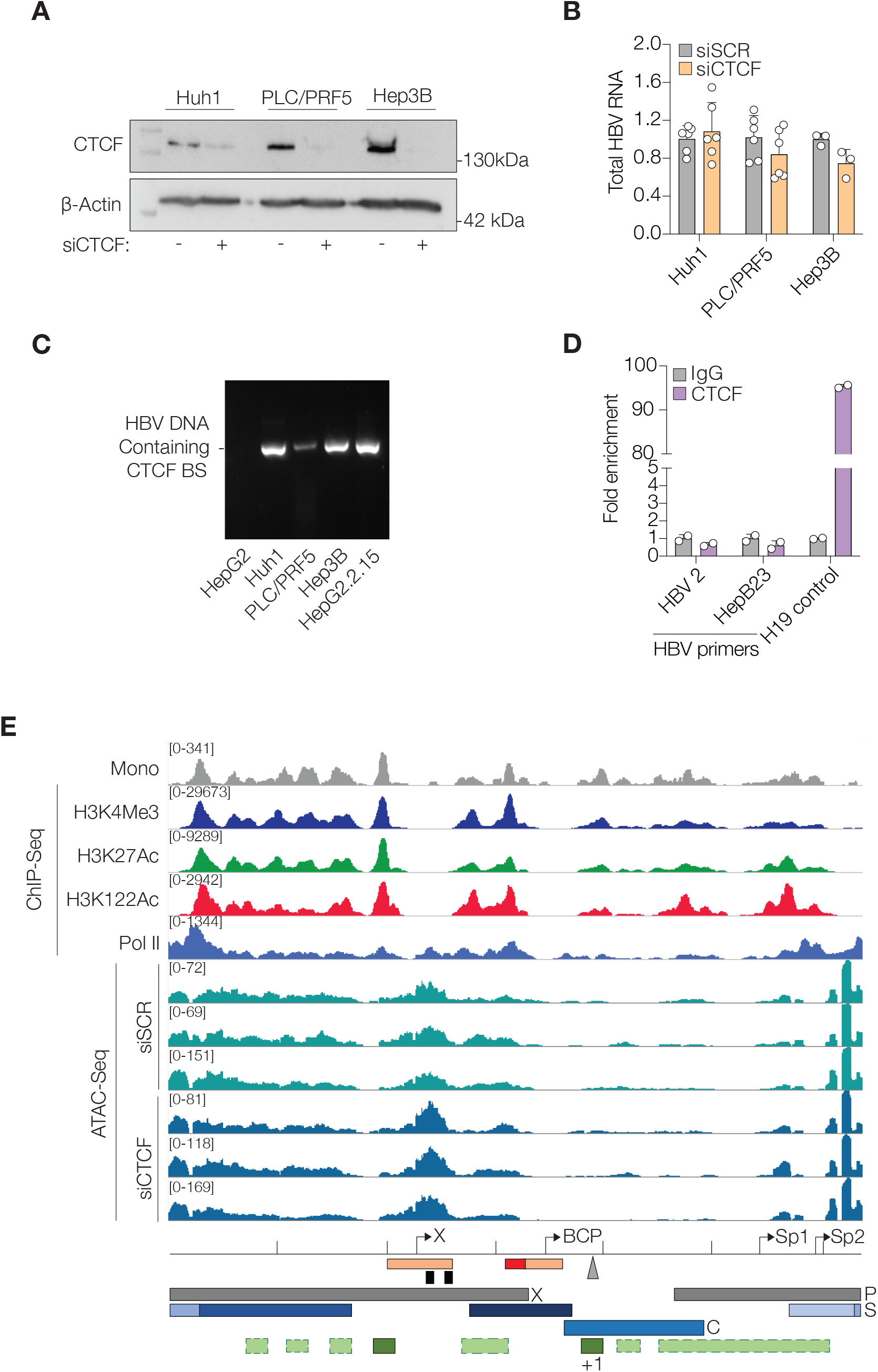
CTCF does not modulate viral transcripts from HBV integrant lines. (A) Huh1, PLC/PRF5 and Hep3B cells were transfected with 25 nM scramble or CTCF siRNA for 72 h. CTCF was detected in the cell lysates by SDS-PAGE followed by immunoblotting using anti-CTCF antibody. Anti-β-actin antibodies were used as a control for loading and a representative image shown. **(B)** RNA was extracted and total HBV RNA measured by RT-qPCR using specific primers. Data is presented as fold change from siSCR controls (mean +/- SD from two independent experiments performed in biological triplicates). **(C)** DNA was extracted from the integrant-bearing lines along with HepG2.2.15 that carry full-length HBV genomes [31] and HepG2 subjected to PCR amplification with HBV primers (HBV2 forward and Hep3B23 reverse) spanning the CTCF binding motifs. Products were separated and visualised by agarose gel electrophoresis. **(D)** CTCF-ChIP assay was performed on PLC/PRF5 cells. Anti-IgG antibodies were used as negative control. Eluted DNA was purified and subjected to qPCR using the indicated HBV primers. Primers for human *H19* were used as a positive control for CTCF binding (Merck Millipore, UK). Data is presented as fold enrichment of CTCF-bound DNA from the corresponding IgG controls. **(E)** Alignment of ATAC-seq analysis of chromatin accessibility in integrated HBV in PLC/PRF5 cells with mononucleosome mapping and ChIP-Seq analysis (H3K4Me3, H3K27Ac, H3K112ac and RNA Pol II) of HBV+ liver tissue as previously reported [27]. Mononucleosome and ChIP-Seq data were downloaded from www.ncbi.nlm.nih.gov/geo (accession no. GSE68402) [27]. ATAC-Seq data alignments are derived from three biological repeats of siSCR or siCTCF-transfected PLC/PRF5 cells. HBV genome features are annotated below including open reading frames, promoters (black arrows), CTCF binding sites (black squares), and polyA+ sites (grey triangle).

CTCF is known to organise chromatin domains and to modify transcription regulatory regions [41]. Tropberger et al., reported differences in the pattern of histone recruitment and modification in HBV DNA isolated from liver tissue compared to experimentally infected hepatocytes [27]. As the clinical samples are likely to contain integrated and episomal genomes [32] we were interested to assess the effect of depleting CTCF on the chromatin accessibility of HBV in the PLC/PRF5 using ATAC-Seq (**Fig.3E**). Comparing our ATAC-Seq data to the available ChIP-Seq data reporting histone modifications in HBV+ liver sample identified distinct peaks in accessible chromatin at the X and SP2 promoters that were flanked by strong histone peaks in the previously published ChIP-Seq study, indicating concordance between the datasets. Notably, depletion of CTCF had no effect on the accessibility of chromatin in the integrated HBV DNA in PLC/PRF5 cells (**Fig.3E**). These data indicate that CTCF is not recruited to integrated HBV DNA in PLC/PRC5 cells and has a negligible role in viral chromatin organisation and transcription control.

### CTCF regulates chromatin accessibility of HBV cccDNA

A previous report characterising the chromatinization of HBV cccDNA reported histones marked with post-translational modifications (PTMs) that are indicative of active transcription including H3K4Me3, H3K27Ac and H3K112Ac [27]. A combination of micrococcal nuclease (MNase) and chromatin immunoprecipitation (ChIP-Seq) experiments identified nucleosomes immediately upstream of the Xp and downstream from the BCP, respectively [27]. These nucleosomes flank an area of chromatin that is either highly dynamically associated with, or devoid of histones that contains Enh I and II. Since the CTCF binding sites in the HBV genome are situated within EnhI, we hypothesised that CTCF may regulate nucleosome phasing and the accessibility of the transcriptional enhancers. To determine whether this is the case, we analysed the accessibility of chromatin in cccDNA established by *de novo* infection of HepG2-NTCP cells with HBV WT or CTCF BS1-2m using ATAC-Seq.

Analysing HBV chromatin structure in *de novo* infected cells revealed an area of open chromatin that spanned Enh I and II and was flanked by phased nucleosomes (**Fig.4**). ATAC-Seq analysis of WT cccDNA revealed a distinct and consistent area of open chromatin that contains the transcriptional enhancer elements Enh I and II, and the Xp and BCP transcriptional start sites (**Fig.4**). Comparison of our ATAC-Seq dataset with previously published MNase and ChIP-Seq data of HBV infected HepG2-NCTP cells suggests that the boundaries of this open area of chromatin align with the two strongly phased nucleosomes upstream of EnhI and at the +1 position upstream of the BCP. Notably, ATAC-Seq analysis of cccDNA derived from *de novo* infection with HBV CTCF BS1-2m revealed a significant reduction in chromatin accessibility (p = 0.036) at the enhancer elements suggesting that CTCF binding with EnhI is important for the specific phasing of nucleosomes to maintain an open chromatin conformation and regulate HBV transcription.

**Figure 4.**
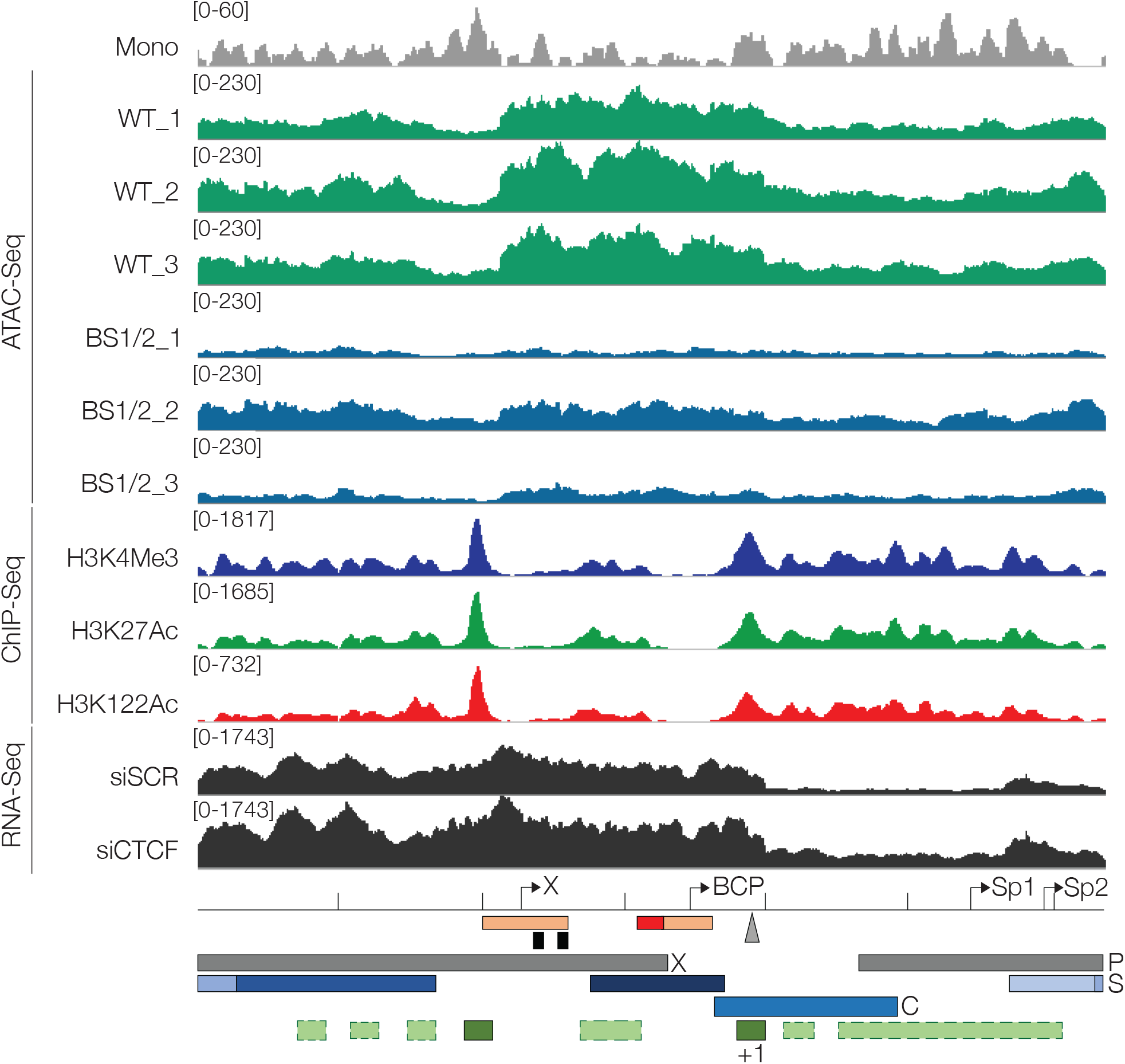
CTCF modifies chromatin accessibility of HBV cccDNA. HepG2-NTCP cells were infected with 400 GE HBV WT or virus harbouring CTCF binding site mutations (HBV BS1-2m) and cultured in media containing 2.5% DMSO. Cells were harvested at 6 dpi and ATAC-Seq performed. Reads were aligned to HBV genome annotated below the figure including open reading frames, promoters (black arrows), CTCF binding sites (black squares), and polyA+ site (grey triangle). Histone phasing in cccDNA as previously described [27] is indicated where light green indicates dynamic association and dark green represents phased nucleosomes up- and downstream of the enhancer elements (orange). The previously described negative regulatory element in HBV EnhII is shown in red. Mononucleosome and ChIP-Seq data (H3K4Me3, H3K27Ac, H3K112ac and RNA Pol II) were downloaded from www.ncbi.nlm.nih.gov/geo (accession no. GSE68402). ATAC-Seq data alignments from three biological repeats of HBV WT- (green) and CTCF BS1-2m- (blue) infected HepG2-NTCP cells are shown and RNA-Seq analysis of HBV mRNAs in siSCR and siCTCF-transfected HepG2-HBV-Epi cells are presented in dark grey.

## DISCUSSION

Previous studies of diverse DNA viruses have highlighted CTCF as an important regulator of virus transcription and transcript processing [42, 43]. While hijacking CTCF is a common mechanism of transcription regulation, there are distinct differences in the way CTCF is utilised between virus families. Herpesviruses including Epstein Barr virus and Kaposi’s sarcoma associated herpesvirus have large (>135kb) episomal genomes that contain multiple CTCF binding sites. These CTCF sites are differentially bound during lytic and latency-associated stages of the virus life cycle, which dictates differential intra-episome chromatin interactions. Differential gene activation in lytic and latent phases is a result of CTCF-dependent stabilisation of discrete chromatin domains and loops within the viral episomes [44]. Human papillomavirus (HPV) also recruits CTCF to regulate viral gene expression but in contrast to herpesviruses with larger genomes, HPV episomes are ∼8kb in size and generally contain a single CTCF binding site [43, 45]. This dominant CTCF binding site is positioned 3kb downstream of the viral transcriptional enhancer and regulates HPV enhancer activity via the stabilisation of a CTCF-YY1 dependent chromatin loop [46]. In this context, CTCF functions as a repressor of HPV transcription in the early stages of the virus life cycle. Derepression in later stages of the life cycle is a result of chromatin loop disruption and activation of the transcriptional enhancer. We previously demonstrated that CTCF functions as a transcriptional repressor of HBV cccDNA [30]. We identified two adjacent CTCF binding sites situated in the viral enhancer I and the downstream transcriptional start site at Xp, respectively. Abrogation of CTCF binding to either one or both CTCF binding sites enhanced transcription in cells transfected with HBV minicircles.

In the present study we extended our analysis of CTCF-dependent HBV transcription repression and analysed the viral transcriptome by next generation sequencing following CTCF depletion in hepatoma cells harbouring cccDNA-like HBV episomes (HepG2-HBV-Epi). These data confirm our earlier findings and provide evidence that CTCF depletion alters the activity of HBV promoters since the increase in viral reads occured throughout the HBV genome. Notably, identification of transcript splicing events within our Illumina data set highlighted a significant increase in SP1 spliced transcript while modest, non-significant increases were observed in the other spliced RNAs. PCR-based validation of these findings confirmed a significant increase in all splice events analysed. We noted a similar increase in the ratio of pC/pg RNA to SP1 spliced transcripts following CTCF depletion in *de novo* infected hepatoma cells confirming a role for CTCF in regulating HBV transcript splicing. However, infection with mutant HBV unable to bind CTCF [30] resulted in increased pC/pgRNA and total HBV RNA but with no change in the relative abundance of SP1 transcripts. These findings suggest that CTCF functions to repress HBV transcription via direct association with cccDNA but that the effect on post-transcriptional processing of HBV transcripts is likely to be via modulation of host cell processes.

It is widely accepted that CTCF regulates alternative splicing (AS) of host mRNAs through both direct and indirect mechanisms (reviewed by [47]). Direct binding of CTCF within AS genes can either create a roadblock to RNA pol II which slows elongation rates and promotes exon inclusion in the nascent mRNA [48], or alter RNA pol II progression by modifying chromatin structure [49]. CTCF can play an indirect role in the regulation of AS, partly through its ability to modulate the activity of Poly (ADP)-ribosylase 1 (PARP1) [50]. Inhibition of PARP1 has been shown to alter AS through deregulation of splicing factors including spliceosomal factor 3B subunit 1 (SF3B1) [51]. Here, we show that depletion of CTCF in HBV infected cells results in an increase in the relative abundance of SP1 transcript. Since this effect on the abundance of spliced transcripts is not observed in mutant HBV that is abrogated in CTCF binding, we conclude that CTCF is likely to play an indirect role in the regulation of HBV transcript AS. These findings contrast with our earlier study which analysed CTCF-dependent transcript processing in HPV where we observed a direct effect of CTCF recruitment to HPV episomes on transcript splicing [43].

Our sequencing study revealed large changes in the host transcriptome of CTCF silenced HepG2 cells. We found that silencing CTCF resulted in the upregulation of n=544 genes, which were associated with several pathways that are exploited by HBV. Fatty acid and bile acid metabolism are hepatocyte specific functions and represent the most significantly upregulated pathways and factors within these gene sets that are being actively explored as therapeutic targets for HBV cure (as reviewed in [52]). In contrast, the most significantly downregulated pathways were genes that are under the transcriptional control of Myc. Myc has been reported to interact with HBV at several levels, and previous studies have shown interactions between the viral accessory protein, HBx, and Myc activation and stability [53, 54]. In short, as well as impacting HBV splicing, CTCF is a potent regulator of the host cell transcriptome, and future investigations into the consequences this has on viral pathogenesis is warranted.

The human leukemia virus, HTLV-1, contains a single CTCF binding site that creates a distinct epigenetic boundary between active and repressed chromatin and acts as an enhancer blocker in the viral DNA [55]. Upon insertion of the provirus into the host chromatin, CTCF, bound to the retroviral DNA, forms long-range interactions with the surrounding chromatin [55, 56]. The insertion of an exogenous CTCF binding site in this manner has profound effects on host cell transcription, thought to be a contributing factor to HTLV-1-driven malignancies. However, this does not appear to be a contributing factor in HBV-mediated disruption of host transcription, at least in the cell lines tested in this study. We show that CTCF depletion in a variety of integrated HBV-carrying hepatoma cell lines does not alter HBV transcription, in contrast to our findings that CTCF depletion in cells that maintain HBV increases HBV transcription. Furthermore, ChIP analysis shows that CTCF was not recruited to integrated viral genomes in PLC/PRF5 cells. These findings suggest that while CTCF is recruited to extrachromosomal cccDNA to regulate HBV transcription, integration of the viral DNA is coincident with loss of CTCF binding. This molecular difference may be responsible for the differential chromatin arrangement of episomal vs integrated HBV DNA [27] and consequently, the different transcriptional profiles observed in episomal and integrated transcriptional templates [57].

In contrast to our finding that depletion of CTCF has no effect on the chromatin structure of integrated HBV, abrogation of CTCF binding in HBV cccDNA established by *de novo* infection resulted in a dramatic change in chromatin topology. WT HBV cccDNA reproducibly contained an area of open chromatin, identified by ATAC-Seq, across the viral enhancers (I/II) and BCP. This area of open chromatin is flanked by histones that are enriched in H3K4Me2, H3K27Ac and H3K112Ac [27]. This arrangement of histones with marks of active transcription phased at the boundary of broad open chromatin areas identified by ATAC-Seq is typical of active enhancers [58] and is presumably required for activation of Xp and BCP. However, our data show that abrogation of CTCF binding within EnhI disrupts the phased nucleosomes and distinctive open chromatin peak identified by ATAC-Seq. This disruption is surprisingly associated with enhanced HBV transcription and could be explained by several hypotheses. Firstly, CTCF could function as an enhancer blocker between EnhI and EnhII and/or the BCP. This has been demonstrated in Herpes simplex virus (HSV-1) where loss of CTCF binding adjacent to the viral transcriptional enhancer results in reactivation of lytic gene expression [59-61]. Alternatively, loss of CTCF recruitment to EnhI disrupts the phased nucleosomes within the HBV enhancer region which disrupts binding of repressive transcription factors such as the Maf bZIP transcription factor F (MafF) [62] or sex determining region Y box2 (SOX2) [63]. Thirdly, it is possible that CTCF binding is important to direct nucleosome phasing at the HBV enhancer, which is important for positioning of the +1 nucleosome immediately downstream of the BCP [27]. Promoter proximal nucleosomes are important for assembly of the RNA polymerase II containing pre-initiation complex but can also inhibit transcription elongation, particularly when the proximal edge is positioned 50-100 nucleotides downstream of the transcription promoter element [64]. The combined analysis of nucleosome position and RNA pol II enrichment described by Tropberger *et al* agrees with this hypothesis; the +1 nucleosome proximal to the BCP is positioned 50-100 bp downstream of the transcriptional start site and significant RNAPol II enrichment is observed upstream of this nucleosome, indicating transcriptional pausing [27]. The loss of the open chromatin region in HBV cccDNA following abrogation of CTCF binding could be due to disruption of the phasing of nucleosomes in relation to the transcriptional elements, resulting in uncontrolled transcriptional elongation and the increased abundance of viral transcripts detected in our analysis.

## FIGURE LEGENDS

**Supplementary Figure 1.**
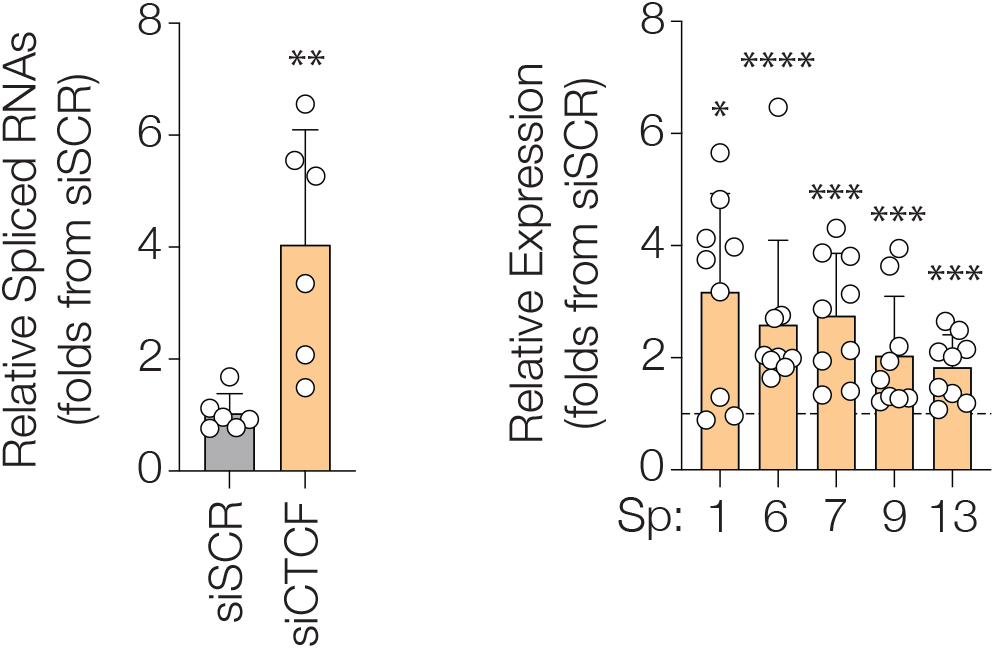
PCR validation of spliced HBV RNAs. The abundance of unique splice junctions was quantified by RT-qPCR using primers detailed in Table 1. Products were Sanger sequenced to confirm amplification of target. Fold expression change was calculated after normalisation to β -actin and shown as fold change compared to siSCR. Data are mean of at least two independent repetitions of experiments performed in triplicates (* p<0.05, *** p<0.001, **** p<0.0001, Mann-Whitney U test).

## ACKNOWLEDGEMENTS

This work was funded by a Medical Research Council (MRC) grant awarded to JLP and JAM (MR/R022011/1). The JAM laboratory is funded by a Wellcome Investigator Award 200838/Z/16/Z, Wellcome Discovery Award 225198/Z/22/Z, Chinese Academy of Medical Sciences Innovation Fund for Medical Science, China (grant number: 2018-I2M-2-002) and AM is supported by the John Black Foundation. The JP lab is funded by MRC research awards (MR/Y001753/1, MR/W031442/1, and MR/T015985/1) and the National Institute for Health and Care Research (NIHR) Birmingham Biomedical Research Centre (BRC). CV was funded by Cancer Research UK (grant No. C17422/A25154). The funders had no role in study design, data collection and interpretation, or the decision to submit the work for publication. We thank Peter Balfe (University of Oxford) for his critical reading of this draft and continued support on this project.

## AUTHOR CONTRIBUTIONS

MOD designed and conducted experiments, analysed data, and co-wrote the paper; CSV designed and conducted experiments, analysed data, and co-wrote the paper; JMH designed experiments, analysed data, and co-wrote the paper; JF analysed data; AM analysed data; RA analysed data; CV analysed data; JLP designed the study, analysed data and co-wrote the paper and JAM designed the study, analysed data and co-wrote the paper.

## REFERENCES

1. Magnius L, Mason WS, Taylor J, Kann M, Glebe D et al. ICTV Virus Taxonomy Profile: Hepadnaviridae. J Gen Virol 2020;101(6):571–572.

2. Ni Y, Lempp FA, Mehrle S, Nkongolo S, Kaufman C et al. Hepatitis B and D viruses exploit sodium taurocholate co-transporting polypeptide for species-specific entry into hepatocytes. Gastroenterology 2014;146(4):1070–1083.

3. Nassal M. HBV cccDNA: viral persistence reservoir and key obstacle for a cure of chronic hepatitis B. Gut 2015;64(12):1972–1984.

4. Lucifora J, Pastor F, Charles E, Pons C, Auclair H et al. Evidence for long-term association of virion-delivered HBV core protein with cccDNA independently of viral protein production. JHEP Rep 2021;3(5):100330.

5. Tong S, Revill P. Overview of hepatitis B viral replication and genetic variability. J Hepatol 2016;64(1 Suppl):S4–S16.

6. Stadelmayer B, Diederichs A, Chapus F, Rivoire M, Neveu G et al. Full-length 5’RACE identifies all major HBV transcripts in HBV-infected hepatocytes and patient serum. J Hepatol 2020;73(1):40–51.

7. Block TM, Chang KM, Guo JT. Prospects for the Global Elimination of Hepatitis B. Annu Rev Virol 2021;8(1):437–458.

8. Tu T, Zhang H, Urban S. Hepatitis B Virus DNA Integration: In Vitro Models for Investigating Viral Pathogenesis and Persistence. Viruses 2021;13(2).

9. Ramirez R, van Buuren N, Gamelin L, Soulette C, May L et al. Targeted Long-Read Sequencing Reveals Comprehensive Architecture, Burden, and Transcriptional Signatures from Hepatitis B Virus-Associated Integrations and Translocations in Hepatocellular Carcinoma Cell Lines. J Virol 2021;95(19):e0029921.

10. van Buuren N, Ramirez R, Soulette C, Suri V, Han D et al. Targeted long-read sequencing reveals clonally expanded HBV-associated chromosomal translocations in patients with chronic hepatitis B. JHEP Rep 2022;4(4):100449.

11. Tu T, Budzinska MA, Shackel NA, Urban S. HBV DNA Integration: Molecular Mechanisms and Clinical Implications. Viruses 2017;9(4).

12. Candotti D, Allain J-P. Biological and clinical significance of hepatitis B virus RNA splicing: an update. Annals of Blood 2016;2:6–6.

13. Lim CS, Sozzi V, Littlejohn M, Yuen LKW, Warner N et al. Quantitative analysis of the splice variants expressed by the major hepatitis B virus genotypes. Microb Genom 2021;7(1).

14. Ng E, Dobrica MO, Harris JM, Wu Y, Tsukuda S et al. An enrichment protocol and analysis pipeline for long read sequencing of the hepatitis B virus transcriptome. J Gen Virol 2023;104(5).

15. Maslac O, Wagner J, Sozzi V, Mason H, Svarovskaia J et al. Secreted hepatitis B virus splice variants differ by HBV genotype and across phases of chronic hepatitis B infection. J Viral Hepat 2022;29(8):604–615.

16. Chen J, Wu M, Wang F, Zhang W, Wang W et al. Hepatitis B virus spliced variants are associated with an impaired response to interferon therapy. Sci Rep 2015;5:16459.

17. Soussan P, Tuveri R, Nalpas B, Garreau F, Zavala F et al. The expression of hepatitis B spliced protein (HBSP) encoded by a spliced hepatitis B virus RNA is associated with viral replication and liver fibrosis. J Hepatol 2003;38(3):343–348.

18. Soussan P, Pol J, Garreau F, Schneider V, Le Pendeven C et al. Expression of defective hepatitis B virus particles derived from singly spliced RNA is related to liver disease. J Infect Dis 2008;198(2):218–225.

19. Bayliss J, Lim L, Thompson AJ, Desmond P, Angus P et al. Hepatitis B virus splicing is enhanced prior to development of hepatocellular carcinoma. J Hepatol 2013;59(5):1022–1028.

20. Soussan P, Garreau F, Zylberberg H, Ferray C, Brechot C et al. In vivo expression of a new hepatitis B virus protein encoded by a spliced RNA. J Clin Invest 2000;105(1):55–60.

21. Sozzi V, McCoullough L, Mason H, Littlejohn M, Revill PA. The in vitro replication phenotype of hepatitis B virus (HBV) splice variant Sp1. Virology 2022;574:65–70.

22. Chen WN, Chen JY, Lin WS, Lin JY, Lin X. Hepatitis B doubly spliced protein, generated by a 2.2 kb doubly spliced hepatitis B virus RNA, is a pleiotropic activator protein mediating its effects via activator protein-1- and CCAAT/enhancer-binding protein-binding sites. J Gen Virol 2010;91(Pt 10):2592–2600.

23. Tsai KN, Chong CL, Chou YC, Huang CC, Wang YL et al. Doubly Spliced RNA of Hepatitis B Virus Suppresses Viral Transcription via TATA-Binding Protein and Induces Stress Granule Assembly. J Virol 2015;89(22):11406–11419.

24. Park GS, Kim HY, Shin HS, Park S, Shin HJ et al. Modulation of hepatitis B virus replication by expression of polymerase-surface fusion protein through splicing: implications for viral persistence. Virus Res 2008;136(1-2):166–174.

25. Revill PA, Chisari FV, Block JM, Dandri M, Gehring AJ et al. A global scientific strategy to cure hepatitis B. Lancet Gastroenterol Hepatol 2019;4(7):545–558.

26. Belloni L, Pollicino T, De Nicola F, Guerrieri F, Raffa G et al. Nuclear HBx binds the HBV minichromosome and modifies the epigenetic regulation of cccDNA function. Proc Natl Acad Sci U S A 2009;106(47):19975–19979.

27. Tropberger P, Mercier A, Robinson M, Zhong W, Ganem DE et al. Mapping of histone modifications in episomal HBV cccDNA uncovers an unusual chromatin organization amenable to epigenetic manipulation. Proc Natl Acad Sci U S A 2015;112(42):E5715–5724.

28. Gehring AJ, Protzer U. Targeting Innate and Adaptive Immune Responses to Cure Chronic HBV Infection. Gastroenterology 2019;156(2):325–337.

29. Pollicino T, Belloni L, Raffa G, Pediconi N, Squadrito G et al. Hepatitis B virus replication is regulated by the acetylation status of hepatitis B virus cccDNA-bound H3 and H4 histones. Gastroenterology 2006;130(3):823–837.

30. D’Arienzo V, Ferguson J, Giraud G, Chapus F, Harris JM et al. The CCCTC-binding factor CTCF represses hepatitis B virus enhancer I and regulates viral transcription. Cell Microbiol 2021;23(2):e13274.

31. Sells MA, Zelent AZ, Shvartsman M, Acs G. Replicative intermediates of hepatitis B virus in HepG2 cells that produce infectious virions. J Virol 1988;62(8):2836–2844.

32. Magri A, Harris JM, D’Arienzo V, Minisini R, Juhling F et al. Inflammatory Gene Expression Associates with Hepatitis B Virus cccDNA-but Not Integrant-Derived Transcripts in HBeAg Negative Disease. Viruses 2022;14(5).

33. Ko C, Chakraborty A, Chou WM, Hasreiter J, Wettengel JM et al. Hepatitis B virus genome recycling and de novo secondary infection events maintain stable cccDNA levels. J Hepatol 2018;69(6):1231–1241.

34. Dobin A, Davis CA, Schlesinger F, Drenkow J, Zaleski C et al. STAR: ultrafast universal RNA-seq aligner. Bioinformatics 2013;29(1):15–21.

35. Liberzon A, Birger C, Thorvaldsdottir H, Ghandi M, Mesirov JP et al. The Molecular Signatures Database (MSigDB) hallmark gene set collection. Cell Syst 2015;1(6):417–425.

36. Turton KL, Meier-Stephenson V, Badmalia MD, Coffin CS, Patel TR. Host Transcription Factors in Hepatitis B Virus RNA Synthesis. Viruses 2020;12(2).

37. Van Damme E, Vanhove J, Severyn B, Verschueren L, Pauwels F. The Hepatitis B Virus Interactome: A Comprehensive Overview. Front Microbiol 2021;12:724877.

38. Lei YC, Hao YH, Zhang ZM, Tian YJ, Wang BJ et al. Inhibition of hepatitis B virus replication by APOBEC3G in vitro and in vivo. World J Gastroenterol 2006;12(28):4492–4497.

39. Wang X, Ao Z, Chen L, Kobinger G, Peng J et al. The cellular antiviral protein APOBEC3G interacts with HIV-1 reverse transcriptase and inhibits its function during viral replication. J Virol 2012;86(7):3777–3786.

40. Zhu YP, Peng ZG, Wu ZY, Li JR, Huang MH et al. Host APOBEC3G protein inhibits HCV replication through direct binding at NS3. PLoS One 2015;10(3):e0121608.

41. Rowley MJ, Corces VG. Organizational principles of 3D genome architecture. Nat Rev Genet 2018;19(12):789–800.

42. Pentland I, Parish JL. Targeting CTCF to Control Virus Gene Expression: A Common Theme amongst Diverse DNA Viruses. Viruses 2015;7(7):3574–3585.

43. Ferguson J, Campos-Leon K, Pentland I, Stockton JD, Gunther T et al. The chromatin insulator CTCF regulates HPV18 transcript splicing and differentiation-dependent late gene expression. PLoS Pathog 2021;17(11):e1010032.

44. Varghese CS, Parish JL, Ferguson J. Lying low-chromatin insulation in persistent DNA virus infection. Curr Opin Virol 2022;55:101257.

45. Paris C, Pentland I, Groves I, Roberts DC, Powis SJ et al. CCCTC-binding factor recruitment to the early region of the human papillomavirus 18 genome regulates viral oncogene expression. J Virol 2015;89(9):4770–4785. 46.

46. Pentland I, Campos-Leon K, Cotic M, Davies KJ, Wood CD et al. Disruption of CTCF-YY1-dependent looping of the human papillomavirus genome activates differentiation-induced viral oncogene transcription. PLoS Biol 2018;16(10):e2005752.

47. Alharbi AB, Schmitz U, Bailey CG, Rasko JEJ. CTCF as a regulator of alternative splicing: new tricks for an old player. Nucleic Acids Res 2021;49(14):7825–7838.

48. Shukla S, Kavak E, Gregory M, Imashimizu M, Shutinoski B et al. CTCF-promoted RNA polymerase II pausing links DNA methylation to splicing. Nature 2011;479(7371):74–79.

49. Ruiz-Velasco M, Kumar M, Lai MC, Bhat P, Solis-Pinson AB et al. CTCF-Mediated Chromatin Loops between Promoter and Gene Body Regulate Alternative Splicing across Individuals. Cell Syst 2017;5(6):628–637 e626.

50. Guastafierro T, Cecchinelli B, Zampieri M, Reale A, Riggio G et al. CCCTC-binding factor activates PARP-1 affecting DNA methylation machinery. J Biol Chem 2008;283(32):21873–21880.

51. Matveeva E, Maiorano J, Zhang Q, Eteleeb AM, Convertini P et al. Involvement of PARP1 in the regulation of alternative splicing. Cell Discov 2016;2:15046.

52. Hyrina A, Burdette D, Song Z, Ramirez R, Okesli-Armlovich A et al. Targeting lipid biosynthesis pathways for hepatitis B virus cure. PLoS One 2022;17(8):e0270273.

53. Terradillos O, Billet O, Renard CA, Levy R, Molina T et al. The hepatitis B virus X gene potentiates c-myc-induced liver oncogenesis in transgenic mice. Oncogene 1997;14(4):395–404.

54. Lee S, Kim W, Ko C, Ryu WS. Hepatitis B virus X protein enhances Myc stability by inhibiting SCF(Skp2) ubiquitin E3 ligase-mediated Myc ubiquitination and contributes to oncogenesis. Oncogene 2016;35(14):1857–1867.

55. Satou Y, Miyazato P, Ishihara K, Yaguchi H, Melamed A et al. The retrovirus HTLV-1 inserts an ectopic CTCF-binding site into the human genome. Proc Natl Acad Sci U S A 2016;113(11):3054–3059.

56. Melamed A, Yaguchi H, Miura M, Witkover A, Fitzgerald TW et al. The human leukemia virus HTLV-1 alters the structure and transcription of host chromatin in cis. Elife 2018;7.

57. Peng B, Jing Z, Zhou Z, Sun Y, Guo G et al. Nonproductive Hepatitis B Virus Covalently Closed Circular DNA Generates HBx-Related Transcripts from the HBx/Enhancer I Region and Acquires Reactivation by Superinfection in Single Cells. J Virol 2023;97(1):e0171722.

58. Arnold M, Stengel KR. Emerging insights into enhancer biology and function. Transcription 2023:1–20.

59. Ertel MK, Cammarata AL, Hron RJ, Neumann DM. CTCF occupation of the herpes simplex virus 1 genome is disrupted at early times postreactivation in a transcription-dependent manner. J Virol 2012;86(23):12741–12759. 60.

60. Washington SD, Edenfield SI, Lieux C, Watson ZL, Taasan SM et al. Depletion of the Insulator Protein CTCF Results in Herpes Simplex Virus 1 Reactivation In Vivo. J Virol 2018;92(11).

61. Amelio AL, McAnany PK, Bloom DC. A chromatin insulator-like element in the herpes simplex virus type 1 latency-associated transcript region binds CCCTC-binding factor and displays enhancer-blocking and silencing activities. J Virol 2006;80(5):2358–2368.

62. Ibrahim MK, Abdelhafez TH, Takeuchi JS, Wakae K, Sugiyama M et al. MafF Is an Antiviral Host Factor That Suppresses Transcription from Hepatitis B Virus Core Promoter. J Virol 2021;95(15):e0076721.

63. Yang H, Mo J, Xiang Q, Zhao P, Song Y et al. SOX2 Represses Hepatitis B Virus Replication by Binding to the Viral EnhII/Cp and Inhibiting the Promoter Activation. Viruses 2020;12(3).

64. Fisher MJ, Luse DS. Promoter proximal nucleosomes attenuate RNA polymerase II transcription through TFIID. J Biol Chem 2023:104928.

